# Mycobiome analyses of critically ill COVID-19 patients

**DOI:** 10.1101/2023.06.13.543274

**Authors:** Danielle Weaver, Sara Gago, Matteo Bassetti, Daniele Roberto Giacobbe, Juergen Prattes, Martin Hoenigl, Florian Reizine, Hélène Guegan, Jean-Pierre Gangneux, Michael John Bromley, Paul Bowyer

## Abstract

**Rationale:** COVID-19-associated pulmonary aspergillosis (CAPA) is a life-threatening complication in patients with severe COVID-19. Previously, acute respiratory distress syndrome in patients with COVID-19 has been associated with lung fungal dysbiosis, evidenced by reduced microbial diversity and *Candida* colonisation. Increased fungal burden in the lungs of critically ill COVID-19 patients is linked to prolonged mechanical ventilation and increased mortality. However, specific mycobiome signatures associated with severe COVID-19 in the context of survival and antifungal drug prophylaxis have not yet been determined and such knowledge could have an important impact on treatment.

**Objectives:** To understand the composition of the respiratory mycobiome in critically ill COVID-19 patients with and without CAPA and the impact of antifungal use in patient outcome.

**Methods:** We performed a multi-national study of 39 COVID-19 patients in intensive care units (ICU) with and without CAPA. Respiratory mycobiome was profiled using ITS1 sequencing and *Aspergillus fumigatus* burden was further validated using qPCR. Fungal communities were investigated using alpha diversity, beta diversity, taxa predominance and taxa abundances.

**Results:** Respiratory mycobiomes of COVID-19 patients were dominated by *Candida* and *Aspergillus.* There was no significant association with corticosteroid use or CAPA diagnosis and respiratory fungal communities. Increased *A. fumigatus* burden was associated with mortality and, the use of azoles at ICU admission was linked with an absence of *A. fumigatus*.

**Conclusions:** Our findings suggest that mould-active antifungal treatment at ICU admission may be linked with reduced *A. fumigatus-*associated mortality in severe COVID-19. However, further studies are warranted on this topic.

## Introduction

COVID-19 is a pulmonary disease caused by severe acute respiratory syndrome coronavirus 2. There have been over 700 million confirmed cases of COVID-19 since December 2019 with mortality ∼7 million^1^. Around 5% of patients with COVID-19 require admission into the intensive care unit (ICU)^2,3^ and, 50% of those patients need mechanical ventilation^4^, thus increasing the risk of hospital-acquired pneumonia^5^. The pulmonary microbiome and its associations with disease outcomes in COVID-19 patients has been explored since the beginning of the pandemic^6–8^. However, our knowledge on the role of fungi in the pathophysiology of COVID-19 is limited. Specifically, the association between respiratory mycobiome composition and patient outcome, and the interplay of antifungal use, is yet to be investigated.

Mycobiome sequencing of the upper respiratory tract (nasopharyngeal swabs) suggests COVID-19 infection significantly reduces fungal diversity, with a higher abundance of *Alternaria* and *Cladosporium* spp., and a lower abundance of other taxa including *Candida* and *Aspergillus*^9^. In the lower respiratory tract (tracheal aspirates), bacterial and fungal microbiome analyses of patients with severe COVID-19 have shown changes over time that might be linked to antimicrobial pressure^10^. A variety of respiratory mycobiome clusters were identified, including those dominated by *Candida* and *Cladosporium.* Using 18S qPCR in bronchoalveolar lavage (BAL) samples, it has been reported that critically ill COVID-19 patients with high fungal burdens are less likely to be liberated from mechanical ventilation^11^. However, the taxa responsible for this outcome remains unclear. Lastly, mycobiome sequencing of BAL found COVID-19 patients with acute respiratory distress syndrome (ARDS) to be associated with reduced fungal diversity and an increase in *Candida* colonisation^12^. In patients without *Candida* colonisation, an increased abundance of an unclassified *Ascomycota* species was identified.

COVID-19-associated pulmonary aspergillosis (CAPA) is an important complication of COVID-19, mainly described in critically ill patients. Multicentre cohort studies of CAPA conducted in the ICU setting report incidence rates varying between 10-15%^14–16^. Nevertheless, mortality rates in patients with CAPA were double that observed in critically ill COVID-19 patients without CAPA^17,18^. Airway epithelial cell damage due to viral replication and COVID-19 associated downregulation of interferon γ signalling pathway, aberrant immune responses due to ARDS, corticosteroids, azithromycin, or the use of immunomodulators have been linked with susceptibility to CAPA^19–22^. With a view to investigating the impact of the respiratory mycobiome in the outcome of COVID-19, we performed a multi-national mycobiome analysis of 39 respiratory samples from critically ill COVID-19 patients with and without CAPA.

## Methods

### Study design, participants, and sample collection

This study was based on a multinational retrospective study on the prevalence of invasive pulmonary aspergillosis in critically ill COVID-19 patients in ICUs during 2020^15^. Inclusion criteria consisted of: PCR confirmed COVID-19 infection, bronchoscopy or tracheal aspiration performed during routine clinical investigations, and chest imaging available seven days before or after respiratory samples were collected. Patients less than 18 years of age were excluded. Respiratory samples not passing quality control (described in Supplemental methods) were also excluded. Respiratory specimens obtained at ICU admission or during ICU stay were collected, aliquoted (at least 1ml) and stored at -80 °C. Criteria for defining aspergillosis were according to previous guidelines^23^ with the following modifications: COVID-19 requiring ICU admission was included as an additional host factor, tracheal aspirates were equated to BAL fluid for microbiological tests, and serum and BAL GM was added as entry criterion. For a summary of full patient demographics, see Table E1.

IRB approval was obtained at each participating center: Medical University of Graz EC #32-296 ex 19/20. University of Genoa Liguria Region Ethics Committee registry number 163/2020. Rennes Teaching Hospital N ° 16-117.

### Sample processing

BAL DNA was extracted using a cetyltrimethylammonium bromide (CTAB) method^24^. Full details on sample processing are provided in the online supplement. Briefly, for mycobiome analysis, the ITS1 region was amplified using Nextera XT compatible versions of ITS1^25^ and ITS2degen (a degenerate version of ITS2 primer) (see Table 1). The presence of *A. fumigatus* in respiratory samples was validated using a TaqMan probe assay targeting the ITS1 region^26^.

### Data analysis

Paired end reads were subject to quality trimming at Q30 and a minimum length filter of 75 nucleotides using bbduk^27^ (BBMap v38.22). Primer sequences were removed using Cutadapt^28^ (v1.18). Reads were mapped to UNITE database using bowtie2^29^ (v2.3.5.1). Count data was further processed in R (v4.1.3) using the following packages: phyloseq^30^ v1.38.0, vegan^31^ v2.5-7, DESeq2^32^ v1.34.0, stringr^33^ v1.4.0, ggplot2^34^ v3.3.5 and tidyr^35^ v1.2.0.

Abundances were standardised to the median sequencing depth. Extremely low abundance taxa were removed by only retaining those occurring > 0.2% in any sample. DESeq2 was used to identify significantly differentially abundant taxa (adjusted *p* value < 0.05 and basemean > 500). Differences in diversity (Shannon, Chao1 and observed OTUs) were assessed using pairwise Wilcoxon rank sum tests. PERMANOVA test was used to assess differences in Bray-Curtis ordination.

## Results

### Patient cohort

The respiratory mycobiome of 91 critically ill COVID-19 patients in the intensive care unit (ICU) was analysed using internal transcribed spacer 1 (ITS1) amplicon sequencing of BAL. Samples from 39 patients harboured significant fungal communities which passed quality control (See Data analysis section in supplementary methods). Table 1 describes demographic and clinical characteristics of the 39 patients maintained in the mycobiome analysis, stratified by CAPA diagnosis. Six patients had CAPA (15%). Patients were from Genoa, Graz and Rennes (64%, 26% and 10%, respectively). Mean age of those with and without CAPA was 61 and 64, respectively. Most patients received systemic corticosteroids (83% of those with CAPA and 73% of those without CAPA). No patients diagnosed with CAPA received azole treatment at the time of ICU admission. Mortality at end of follow-up in patients with CAPA was 33% (2/6) while mortality in patients without CAPA was also 33% (11/33).

**Table 1.** Clinical and demographic characteristics of patients with and without probable CAPA diagnosis.

| Variable | Patients with CAPA (N=6) | Patients without CAPA (N=33) |
| --- | --- | --- |
| Age |  |  |
| Mean (SD) | 61 (3.4) | 64 (7.9) |
| valid (missing) | 6 (0) | 32 (1) |
| Sex |  |  |
| male | 100% (6) | 79% (26) |
| female | 0% (0) | 18% (6) |
| missing | 0% (0) | 3% (1) |
| Ethnicity |  |  |
| Caucasian | 100% (6) | 91% (30) |
| other |  | 6.1% (2) |
| missing | 0% (0) | 3% (1) |
| BMI > 30 |  |  |
| yes | 50% (3) | 18% (6) |
| no | 50% (3) | 79% (26) |
| missing | 0% (0) | 3% (1) |
| Smoking |  |  |
| yes | 33% (2) | 12% (4) |
| no | 67% (4) | 85% (28) |
| missing | 0% (0) | 3% (1) |
| Institution |  |  |
| Graz | 33% (2) | 24% (8) |
| Genoa | 17% (1) | 73% (24) |
| Rennes | 50% (3) | 3% (1) |
| Hematology oncology |  |  |
| yes | 17% (1) | 9.1% (3) |
| no | 83% (5) | 88% (29) |
| missing | 0% (0) | 3% (1) |
| Solid organ transplant |  |  |
| yes | 17% (1) | 3% (1) |
| no | 83% (5) | 94% (31) |
| missing | 0% (0) | 3% (1) |
| Cardiovascular disease |  |  |
| yes | 67% (4) | 45% (15) |
| no | 33% (2) | 52% (17) |
| missing | 0% (0) | 3% (1) |
| Pulmonary disease |  |  |
| yes | 33% (2) | 24% (8) |
| no | 67% (4) | 73% (24) |
| missing | 0% (0) | 3% (1) |
| Diabetes Mellitus |  |  |
| yes | 17% (1) | 9.1% (3) |
| no | 83% (5) | 88% (29) |
| missing | 0% (0) | 3% (1) |
| Corticosteroids |  |  |
| yes | 83% (5) | 73% (24) |
| no | 17% (1) | 24% (8) |
| missing | 0% (0) | 3% (1) |
| Tocilizumab |  |  |
| yes | 17% (1) | 6.1% (2) |
| no | 83% (5) | 91% (30) |
| missing | 0% (0) | 3% (1) |
| Azithromycin |  |  |
| yes | 33% (2) | 33% (11) |
| no | 67% (4) | 61% (20) |
| missing | 0% (0) | 6.1% (2) |
| Azole treatment at ICU admission |  |  |
| yes | 0% (0) | 15% (5) |
| no | 100% (6) | 6.1% (2) |
| missing | 0% (0) | 79% (26) |
| Life Support |  |  |
| mechanical | 100% (6) | 70% (23) |
| ECMO |  | 3% (1) |
| noninvasive |  | 6.1% (2) |
| mechanical & noninvasive |  | 12% (4) |
| none |  | 6.1% (2) |
| missing | 0% (0) | 3% (1) |
| Duration ICU (days) |  |  |
| Mean (SD) | 27 (11) | 32 (28) |
| valid (missing) | 6 (0) | 32 (1) |
| Palliative (Day 28 or 32) |  |  |
| yes | 67% (4) | 48% (16) |
| no | 33% (2) | 48% (16) |
| missing | 0% (0) | 3% (1) |
| Survival (at end of follow up) |  |  |
| yes | 33% (2) | 33% (11) |
| no | 67% (4) | 64% (21) |
| missing | 0% (0) | 3% (1) |
| Days from ICU admission to CAPA |  |  |
| Mean (SD) | 6.2 (3.5) | - |
| BAL GM (ODI > 1) |  |  |
| positive | 83% (5) | 9.1% (3) |
| negative | 17% (1) | 82% (27) |
| missing | 0% (0) | 9.1% (3) |
| BAL PCR |  |  |
| positive | 50% (3) | 0% (0) |
| negative | 0% (0) | 58% (19) |
| missing | 50% (3) | 42% (14) |
| BAL culture |  |  |
| positive | 50% (3) | 0% (0) |
| negative | 50% (3) | 97% (32) |
| missing | 0% (0) | 3% (1) |
| BAL LFD |  |  |
| positive | 17% (1) | 6.1% (2) |
| negative | 0% (0) | 9.1% (3) |
| missing | 83% (5) | 85% (28) |
| Tracheal Aspirate GM |  |  |
| positive | 0% (0) | 3% (1) |
| negative | 0% (0) | 3% (1) |
| missing | 100% (6) | 94% (31) |
| Tracheal Aspirate PCR |  |  |
| positive | 33% (2) | 0% (0) |
| negative | 17% (1) | 21% (7) |
| missing | 50% (3) | 79% (26) |
| Tracheal Aspirate Culture |  |  |
| positive | 17% (1) | 0% (0) |
| negative | 33% (2) | 3% (1) |
| missing | 50% (3) | 97% (32) |
| Serum GM (> 0. 5) |  |  |
| positive | 33% (2) | 0% (0) |
| negative | 50% (3) | 42% (14) |
| missing | 17% (1) | 58% (19) |
| Bronchial Aspirate culture |  |  |
| positive | 17% (1) | 0% (0) |
| negative | 33% (2) | 21% (7) |
| missing | 50% (3) | 79% (26) |

CAPA; COVID-19 associated pulmonary aspergillosis. BMI; body mass index. ICU; intensive care unit. ECMO; extracorporeal membrane oxygenation. BAL; bronchoalveolar lavage. GM; galactomannan. LFD; lateral flow device.

### *Candida* and *Aspergillus spp.* dominate respiratory mycobiomes in critically ill COVID-19 patients

Median read counts per sample for the 39 samples that passed quality control was 57,913 (range 6,970 – 342,640). There were 36 Genera in total and a median of 5 Genera per sample (range 1 – 22). There was no significant clustering between sample batches (Fig. E1), suggesting that sample processing had no impact on mycobiome communities. Mycobiomes predominantly consisted of *Candida* and *Aspergillus* (Fig. 1A). *Candida albicans*, *Aspergillus fumigatus* and *Candida parapsilosis* were the most abundant species (Fig. E2A-B).

**Fig. 1.**
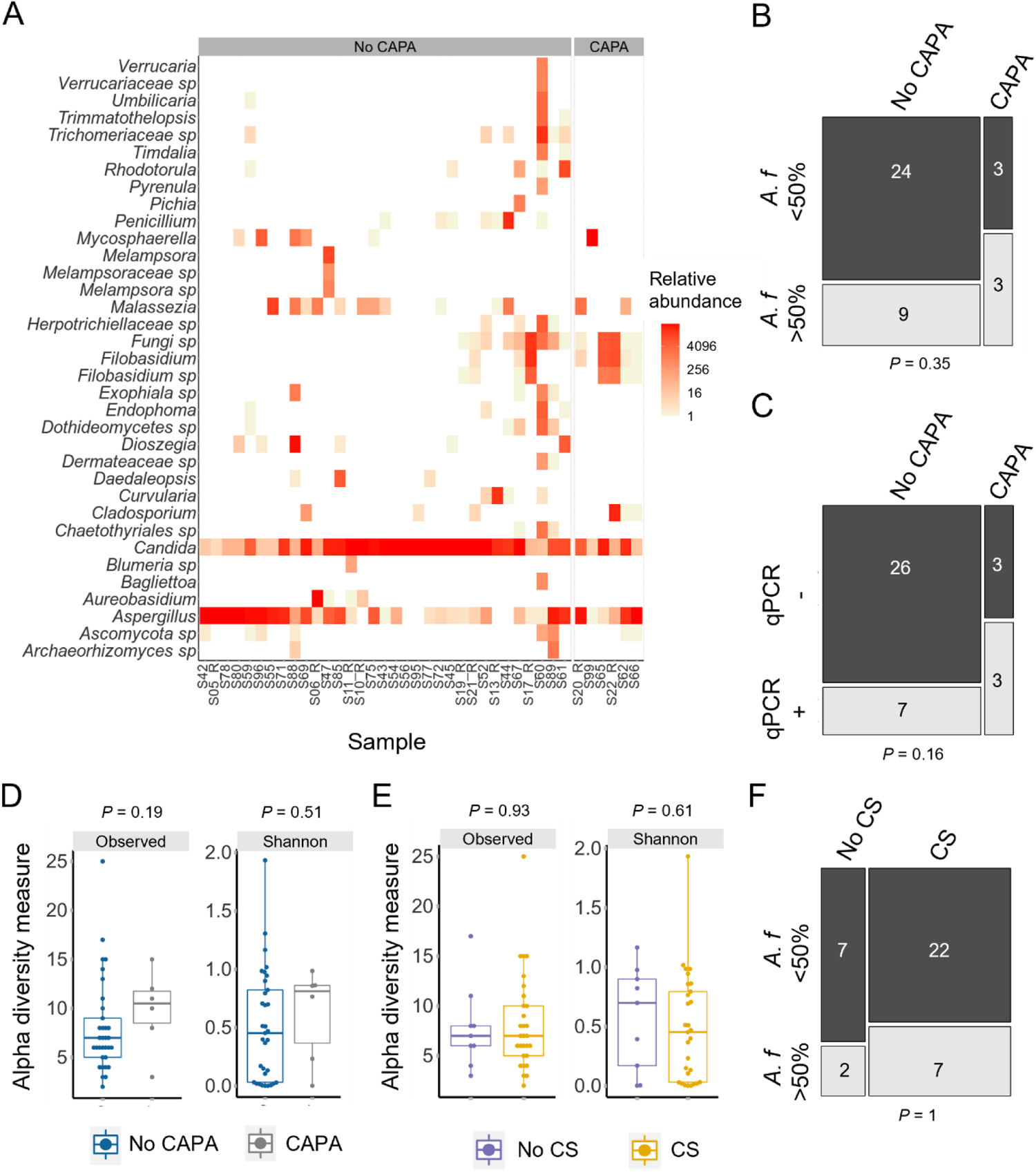
*Aspergillus* and *Candida spp.* dominate the respiratory mycobiome in critically ill COVID-19 patients. (A) *Aspergillus* and *Candida* were the main genera observed in the lungs from COVID-19 patients included in the study (n = 39). Samples are grouped based on CAPA status. (B) *A. fumigatus w*as the predominant species in the mycobiomes of 50% of patients with COVID-19 associated pulmonary aspergillosis (CAPA), compared to 27% of those without CAPA. (C) Fifty percent of CAPA patients were *A. fumigatus-*positive by species specific qPCR, compared to 21% of those without CAPA. (D) Alpha diversity measures (Observed OTUs and Shannon) trended towards higher diversity in CAPA patients (E) Corticosteroid treatment caused no apparent effect on alpha diversity as measured by Observed OTUs or Shannon diversity. (F) *A. fumigatus w*as the predominant species in the mycobiomes of 24% of patients receiving corticosteroids, compared to 22% of those without corticosteroids. Hypothesis testing was applied using Wilcoxon Rank Sum tests (D,E) or Fisher’s exact tests (B,C,F). CS: Corticosteroids. Boxplot data represent median and interquartile range.

### No significant correlation is found between mycobiome communities and CAPA status or corticosteroid use

In our study, higher median *A. fumigatus* levels were observed in CAPA when assessed by read count (∼16,700 vs. 35) and *A. fumigatus* specific qPCR (0.3 vs. 0 genome equivalents) (Fig. E3A-B), but there was overlap between the patient groups and statistical significance was not reached. *A. fumigatus* burden in patients without CAPA was varied, some patients had little to no *A. fumigatus* and others contained particularly high burden (Fig. 1A, Fig. E3A-B). As *A. fumigatus* levels appeared to be bimodal, we dichotomised these data into two groups to assess either the predominance of *A. fumigatus* (present at over 50% of a mycobiome sample) or high *A. fumigatus* burden (qPCR positive at over 0.1 haploid genome equivalents (HGE)). Analysing the data in this manner also found no significant difference between *A. fumigatus* predominance (Fig. 1B) or high burden (Fig. 1C) in patients with and without CAPA. Furthermore, species differential abundance analysis did not find significant differences between *A. fumigatus* levels based on CAPA status. Instead, *Cladosporium delicatulum, Mycosphaerella tassiana* and *Filobasidium magnum* were found to be at higher abundance in CAPA patients (Fig. E6). In addition, probable CAPA patients had higher median α diversity, however, this difference was not significant, and no significant effect on β diversity was observed (Fig. 1D, Fig. E3C).

Corticosteroid use resulted in lower median α diversity (Shannon only), however, this difference was not significant (Fig. 1E). Median levels of *A. fumigatus* were lower in the corticosteroid treated group as assessed by sequencing (8 vs. ∼5,800 reads) and, qPCR (0 vs. 0.1 genome equivalents) (Fig.E3D-E). However, patients on corticosteroids were highly heterogenous in terms of *A. fumigatus* abundance. There was no significant difference between *A. fumigatus* predominance (Fig. 1F) or high burden (Fig. E3F) in patients with and without corticosteroid treatment. Species differential abundance analysis found no significant differences between corticosteroid usage. In addition, corticosteroid use had no significant impact on β diversity (Fig. E3G).

### Increased *A. fumigatus* burden is associated with mortality

The mycobiome of surviving individuals showed a trend towards higher median α diversity (Fig. 2A). Grouped mean abundances indicated a lower level of *Aspergillus* was present upon survival (Fig.2B). At the individual sample level, many mycobiomes of non-surviving patients were predominated by *Aspergillus* (Fig. E4). Species predominance analysis suggested that this difference was due to *A. fumigatus,* with 32% (8/25) of non-surviving patients’ mycobiomes being dominated by this species compared to only 8% (1/13) of patients which survived (Fig. 2C-D). Furthermore, mycobiome differential abundance analysis found *A. fumigatus* and *C. albicans* to be significantly less abundant upon survival, with log fold change values of -4.3 and -4.6, respectively (*padj* < 0.05) (Fig.2E). Quantitative PCR data also showed 32% of patients which did not survive displayed a high burden of *A. fumigatus* compared to only 8% of surviving patients (Fig. 2F). All datapoints for *A. fumigatus* relative abundance and qPCR burden are shown in Fig. E5A-B. BAL galactomannan index values were not significantly different between the patient groups, with an outlier in the survival group having a very high index (Fig.E5C).

**Fig. 2.**
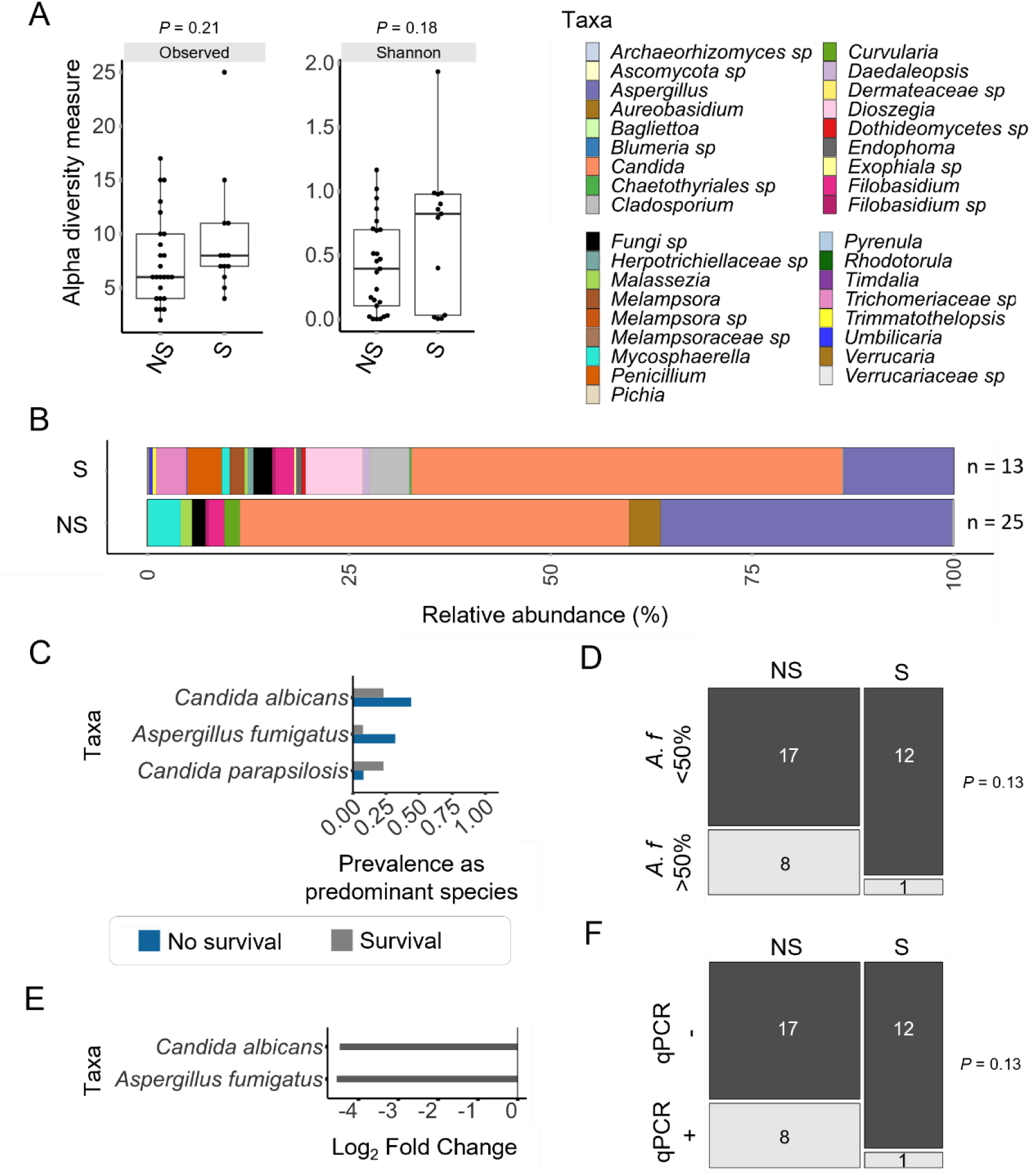
A higher *A. fumigatus* burden is associated with mortality in critically ill COVID-19 patients. (A) Alpha diversity measures (Observed OTUs and Shannon) trended towards higher diversity in critically ill COVID-19 patients which survived. Data represent median and interquartile range. (B) At the genus level, pooled relative abundance mycobiome data from patients which survived (n =13) indicated a lower proportion of *Aspergillus*, and an apparent increase in the number of observed taxa overall. (C) *C. albicans* and *A. fumigatus* were prevalent as the predominant species in a mycobiome in more patients which did not survive (blue, prevalence 0.44 and 0.32, respectively) than those that did survive (grey, prevalence 0.32 and 0.08, respectively). Taxa were counted if present at over 50% of total counts, and only taxa found in at least 10% of samples of either group are shown. (D) *A. fumigatus* was the predominant species in 8/25 (32%) patients which did not survive, compared to 1/13 (8%) of patients which did survive. (E) Analysis using DESeq2 identified *A. fumigatus* and *C. albicans* to be at significantly lower abundance in patients which survived. (F) Thirty-two percent of patients which did not survive were *A. fumigatus-*positive by species specific qPCR, compared to 7% of those which survived. Hypothesis testing in was applied using Wilcoxon Rank Sum tests (A), Fisher’s exact tests (D,F), or DESeq2 (E). S: Survival; NS: No survival.

### Azole treatment at intensive care unit admission is associated with reduced A. fumigatus burden in critically ill COVID-19 patients and COVID-19 survival

Use of azole treatment at ICU admission in critically ill COVID-19 patients resulted in significantly reduced α diversity when analysing raw mycobiome data. Upon removal of very rare taxa, there was a trend towards reduced median α diversity upon azole treatment (Fig. 3A). There was a lack of *Aspergillus* in the mycobiomes of patients with azole treatment, and *Aspergillus* was present at a considerable relative abundance in ∼38% (3/8) patients receiving azoles (Fig. 3B). At the species level, *A. fumigatus* was the predominant species in 25% patients without azole treatment, whereas this species was not detectable in patients receiving treatment (Fig. 3C-D). Furthermore, differential abundance analysis of mycobiome data found a significant reduction of *A. fumigatus* in patients which received azole treatment (LFC -6.3, *padj* 0.04) (Fig. 3E). This analysis also found *Candida albicans, Candida parapsilosis* and *Candida tropicalis* had significantly higher abundance in COVID-19 patients receiving azole treatment (LFC 5.3, 9.4 and 14.5, respectively). Quantitative PCR (qPCR) data found 38% of patients which did not receive azoles displayed a high burden of *A. fumigatus* compared to no patients on azole treatment (Fig. 2F). Datapoints for *A. fumigatus* relative abundance and qPCR burden with and without azole treatment at ICU admission are shown in Fig. E5D-E. BAL galactomannan index was not statistically different between patients with or without azole treatment, however, all individuals receiving treatment were GM negative whereas only half of the patients without treatment were GM negative (Fig. 3G).

**Fig. 3.**
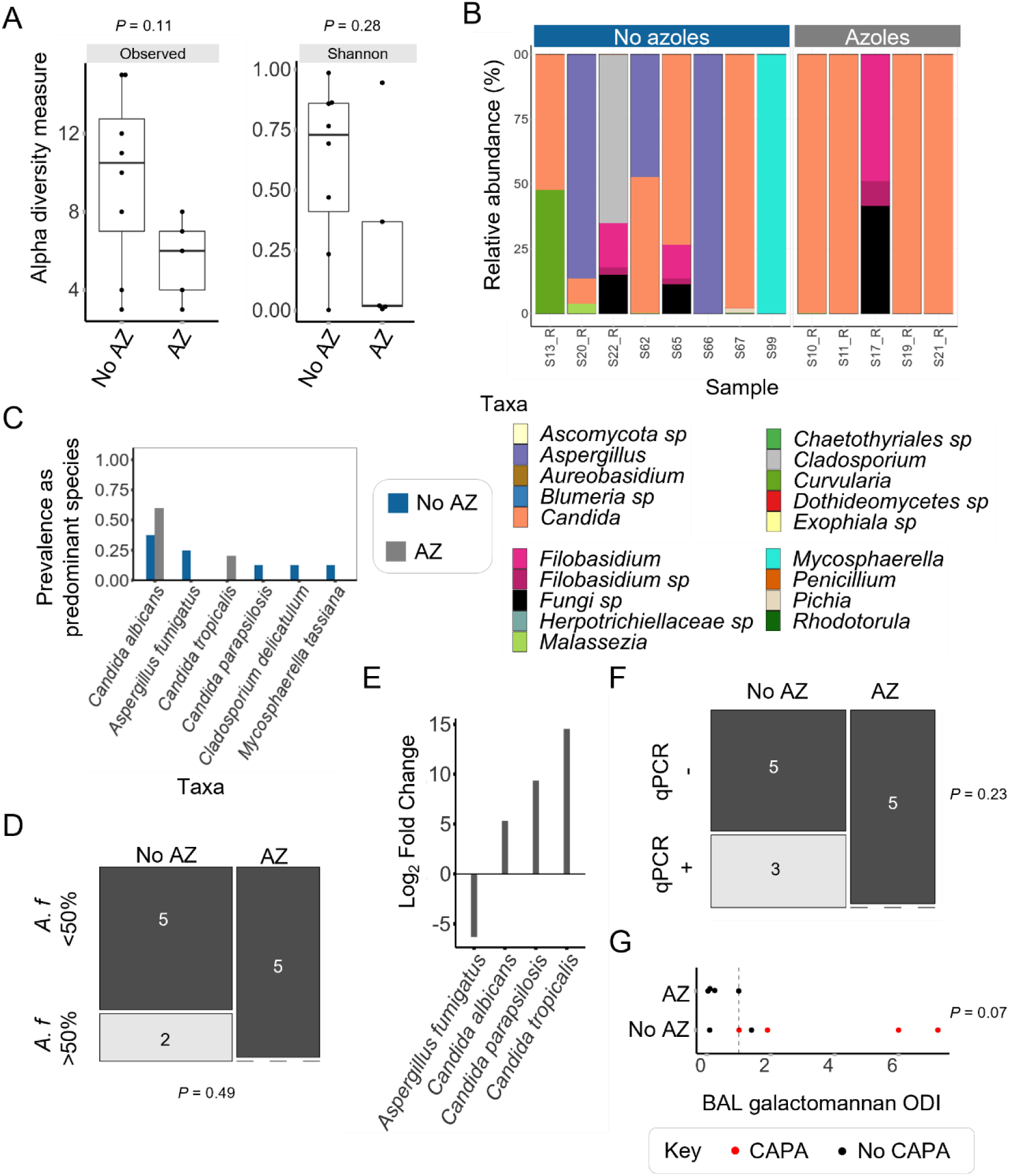
*A. fumigatus* is associated with the absence of azole treatment at intensive care unit admission in critically ill COVID-19 patients. (A) Alpha diversity measures (Observed OTUs and Shannon) trended higher diversity in COVID-19 patients without azoles at ICU admission (AZ). (B) At the genus level, BAL ITS1 mycobiomes from COVID-19 patients which received azoles (n =5) display an absence of *Aspergillus,* whereas 3/8 patients not receiving azoles harboured *Aspergillus.* (C) *C. albicans* was prevalent as the predominant species of a mycobiome in more patients which received azoles (grey, prevalence 0.6) than those which did not receive azoles (blue, prevalence 0.38). *A. fumigatus* was prevalent as the predominant species in 25% of patients receiving azoles. *A. fumigatus* was not prevalent in any patients which received azoles. Taxa were counted if present at over 50% of total counts, and only taxa found in at least 10% of samples of either group are shown. (D) *A. fumigatus* was the predominant species in 2/8 (25%) patients which did not survive, compared to none of the patients which did survive. (E) Analysis using DESeq2 identified *A. fumigatus* to be at significantly lower abundance in patients which received azoles. *C. albicans, C. parapsilosis* and C. *tropicalis* were all at significantly higher abundance in patients receiving azoles. (F) Thirty eight percent (3/8) of patients which did not receive azoles were *A. fumigatus-*positive by species specific qPCR, compared to no patients which did receive azoles. (G) All patients receiving azoles were BAL galactomannan negative (ODI 1 or lower). Two thirds (4/6) of patients not receiving antifungals were galactomannan positive. Hypothesis testing in was applied using Wilcoxon Rank Sum tests (A, G), Fisher’s exact tests (D,F), or DESeq2 (E). AZ: Azoles; No AZ: No azoles; ODI: Optical density index.

Our findings suggest an association between *A. fumigatus* abundance and mortality in critically ill COVID-19 patients, and that azole treatment at ICU significantly reduces *A. fumigatus* levels. Therefore, we combined these factors to assess the association between *A. fumigatus* and survival outcomes depending on the presence or absence of azole treatment. It was apparent that *Aspergillus* was associated with mortality only in COVID-19 patients who had not received azole treatment (Fig. 4A). Presence of *A. fumigatus* in only those patients that did not survive or receive azole treatment was confirmed by qPCR (Fig. 4B). Therefore, these findings suggest azole treatment at ICU admission may have been protective against *A. fumigatus-*associated mortality in this severe COVID-19 patient cohort.

**Fig. 4.**
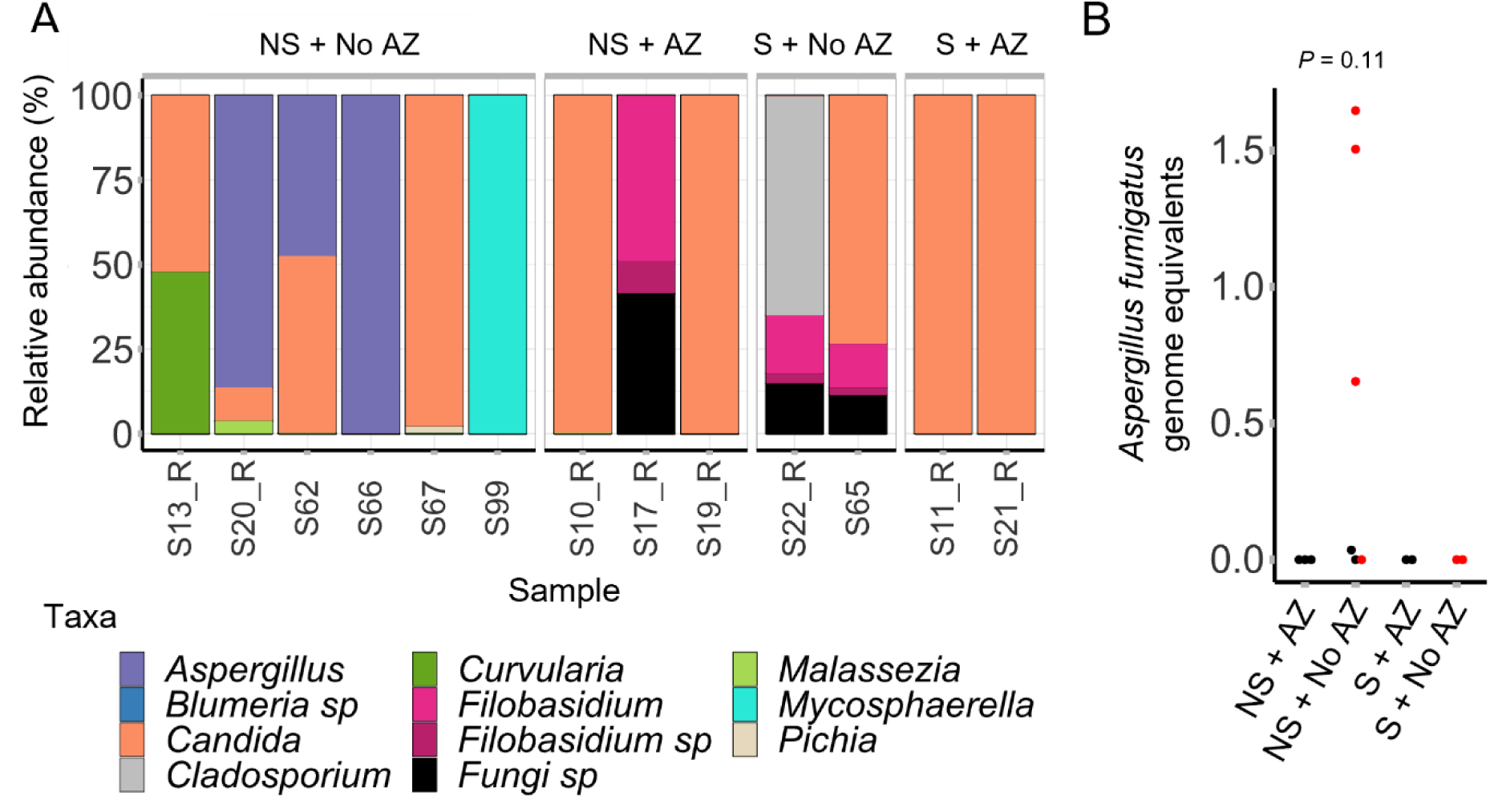
*A. fumigatus* is associated with mortality in patients with COVID-19 who have not received azole treatment at intensive care unit admission. (A) When combining the use of azoles at ICU and survival outcomes, *Aspergillus* was found in the BAL ITS1 mycobiomes from 50% of patients which did not receive azoles or survive (3/6). No *Aspergillus* was present in samples from any other groups. (B) *A. fumigatus* levels (measured by qPCR) did not differ significantly between groups (Wilcoxon Rank Sum test); however, *A. fumigatus* burden was observed only in the patient group which did not survive or receive azole treatment at ICU admission. AZ: Azoles. S: Survival. NS: No survival.

## Discussion

This multinational study found higher *Aspergillus fumigatus* levels in critically ill COVID-19 patients were associated with increased mortality. In addition, the association of *A. fumigatus* with mortality was found only in patients who did not receive azole treatment at ICU admission, suggesting that the use of prophylactic mould-active antifungals in severe COVID-19 patients is potentially valuable for the reduction of *A. fumigatus*-associated mortality in this cohort.

The respiratory mycobiomes of critically ill COVID-19 patients described here were dominated by *Candida* and *Aspergillus.* Previous studies using similar patient groups have also reported lung fungal communities to be dominated by *Candida*^10,12^. Furthermore, one study identified a significant increase in unidentified Ascomycota spp. in patients without *Candida* colonisation^12^. Due to the reported incidence of CAPA in severe COVID-19, the authors hypothesised that *Aspergillus* could be present in these patients and, although mycobiome samples were mostly negative for *Aspergillus*, the presence of *Aspergillus* was confirmed by PCR in follow up BAL samples in over 20% of patients.

A recent report suggests higher fungal burden in the lung microbiota of patients with proven/probable CAPA^11^. However, considerable overlap in fungal burdens between those with and without CAPA was noted. Our study found CAPA patients had higher median levels of *A. fumigatus* and trended towards higher fungal diversity. However, these findings did not meet statistical significance, which may have been driven by the low sample size (n = 6 in CAPA group). Some patients within the non-CAPA group harboured considerable levels of *A. fumigatus.* It is known that high *Aspergillus* burdens can be found in the lungs of healthy individuals^36^. These observations suggest that if a sufficient level of *Aspergillus* is present in the lung, other factors such as disease susceptibility or strain virulence in the context of CAPA may be more important than burden in the outcome of infection. This study was limited to one sample time point, and it would be interesting to assess how *Aspergillus* burden changes during CAPA or COVID-19 infection.

It has been suggested that corticosteroid treatment increases lung fungal burden (particularly *A. fumigatus*)^13^ and lowers Shannon diversity^37^ in asthma. In contrast, no significant differences were found between mycobiome diversity or taxa abundance in respiratory fungal communities of COPD patients with or without inhaled corticosteroid treatment^38^. Another recent study found that alterations in the airway mycobiome in COPD were not significantly affected by corticosteroid use^39^. Large cohort studies suggest systemic corticosteroids are a risk factor for CAPA^16,19^, however, there are no specific reports of the influence corticosteroid use has on lung fungal communities in COVID-19. In our study, patients receiving corticosteroids displayed lower median levels of *A. fumigatus* and lower fungal diversity compared to those not receiving corticosteroids. However, these differences did not meet statistical significance. As most (∼74%) patients received corticosteroids in this cohort, this may have contributed to low power in these statistical comparisons, warranting further data on this topic.

A high fungal burden has previously been associated with a lower likelihood of release from mechanical ventilation and increased mortality risk in severe COVID-19 patients^11^. However, as this study utilised pan-fungal qPCR to identify burden, there was no indication of the specific fungal taxa responsible for this association. Our findings suggest that higher levels of *A. fumigatus* are associated with increased mortality in severe COVID-19. There are no previous reports on the impact of antifungal use on the respiratory mycobiome in COVID-19 patients. In this study, the use of azoles at ICU admission was associated with an absence of *A. fumigatus* and appeared protective against *A. fumigatus-*associated mortality.

Our study investigated the composition of respiratory fungal communities in critically ill COVID-19 patients with and without CAPA. *Candida* and *Aspergillus* were predominant in the respiratory communities. CAPA diagnosis was associated with higher median *A. fumigatus* level and fungal diversity, and a higher prevalence of *A. fumigatus* was associated with mortality and a lack of azole treatment at ICU admission. Our data suggests that the potential use of prophylactic antifungals (with anti-*Aspergillus* activity) in seriously ill COVID-19 patients is worthy of further consideration for the possible prevention of *A. fumigatus*-associated mortality. However, a limitation of this study is the small number of patients included, particularly the low number of CAPA cases. In addition, incomplete clinical data with respect to azole use reduced the sample sizes for this comparison, which may have resulted in low power for these statistical tests. Therefore, study of a larger cohort would be valuable to improve our understanding of the association between prophylactic azole use, the presence of *A. fumigatus* in the respiratory mycobiome and patient outcome in COVID-19 critical care.

## Acknowledgements

This work was part of the ECMM CAPA initiative under the umbrella of the ECMM NGS Working group. This work was funded by the NIHR Manchester Biomedical Research Centre (NIHR203308). The views expressed are those of the author(s) and not necessarily those of the NIHR of the Department of Health and Social Care.

## Supplementary data

Raw sequence data has been deposited at the NCBI sequence read archive (SRA) under accession number PRJNA905224. Code used for analysis is available at https://github.com/Danweaver1/COVID_respiratory_mycobiome.

**Fig. E1.**
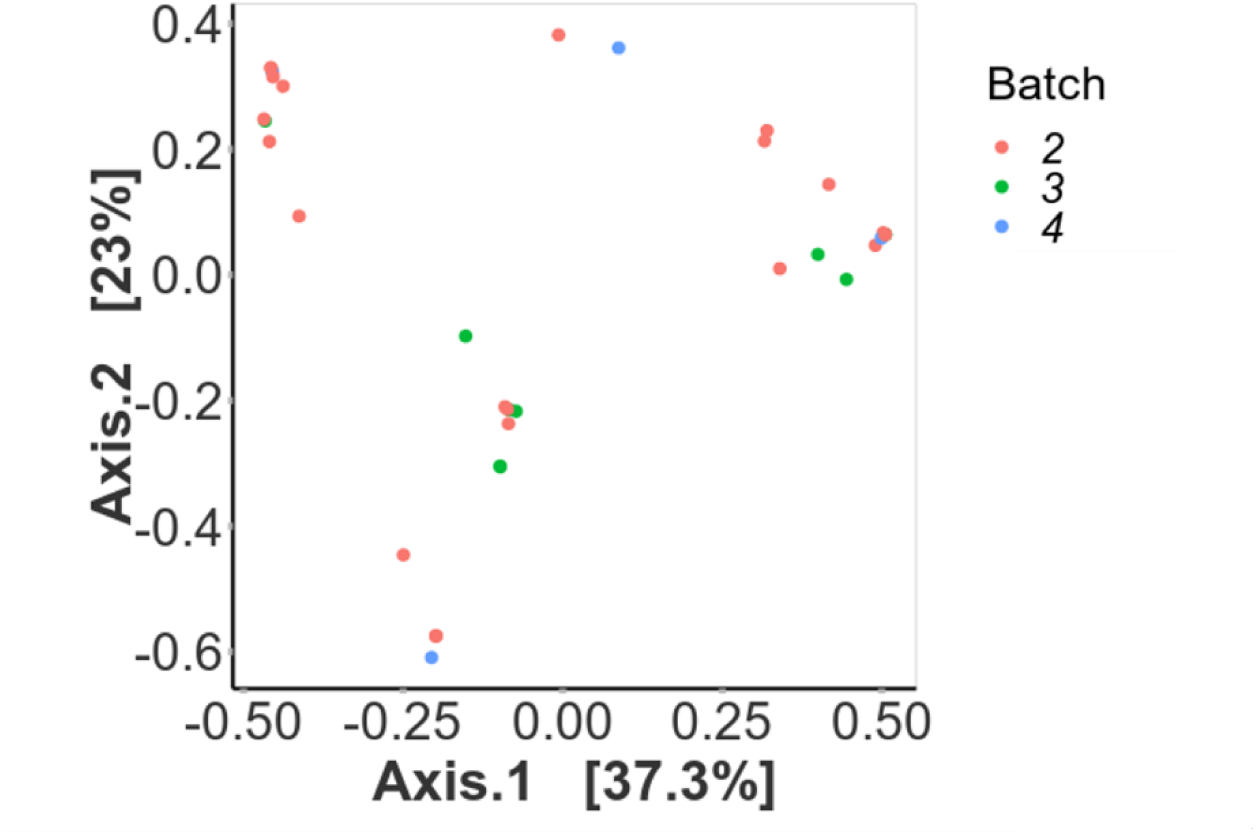
Mycobiome Bray-Curtis ordination by sample batch.

**Fig. E2.**
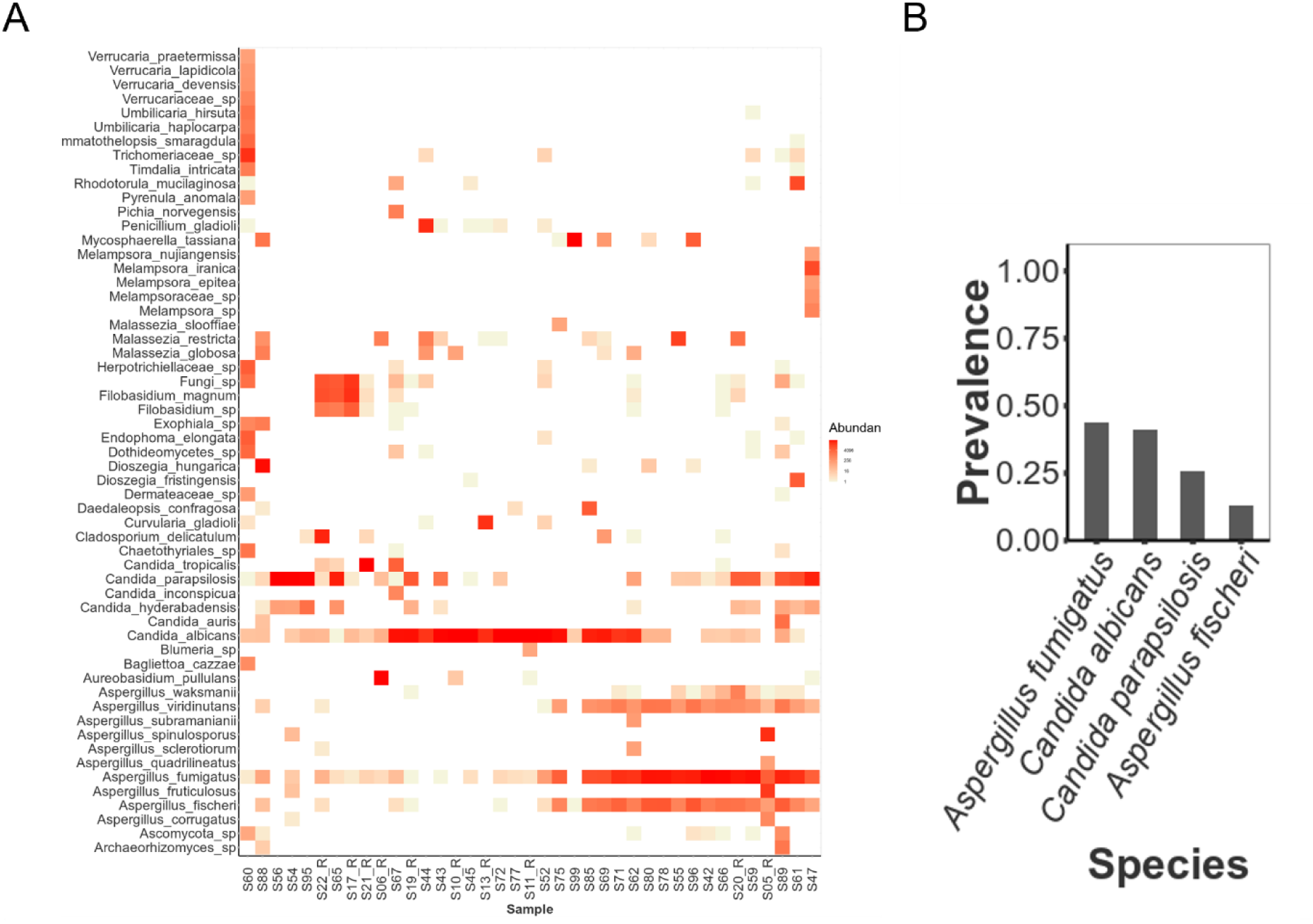
ITS1 respiratory mycobiomes at the species level. (A) Heatmap of fungal species identified in all samples. To remove extremely rare taxa, only those present at > 0.2% in one sample were retained. (B) *A. fumigatus* and *C. albicans* are the most prevalent species. Taxa were counted if present at over 5% in a sample, and only taxa found in at least 10% of samples of either group are shown.

**Fig. E3.**
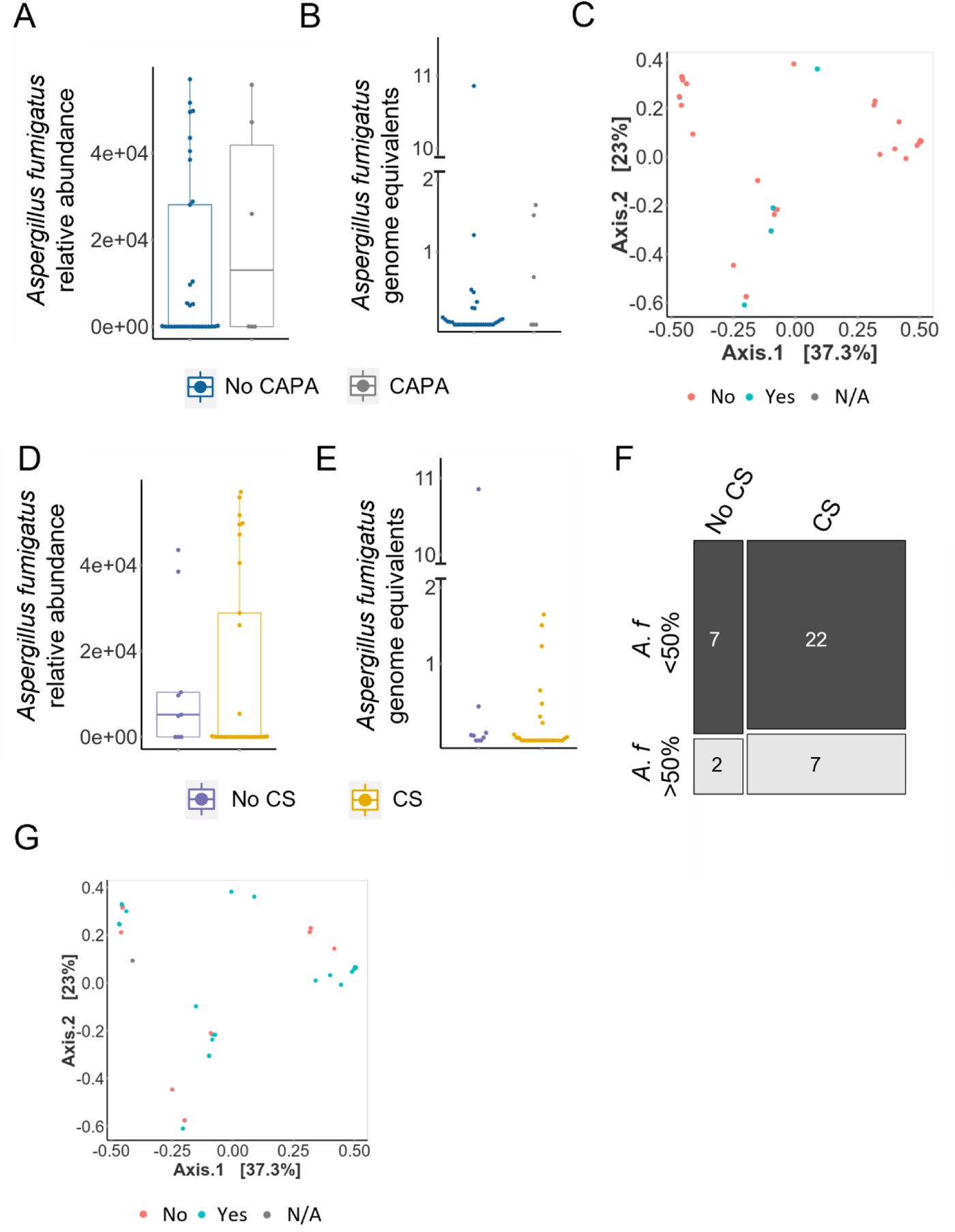
CAPA diagnosis and corticosteroid use had no significant impact on fungal burden, *Aspergillus fumigatus* levels or beta diversity. (A) Relative abundance of *A. fumigatus* in mycobiomes of those with and without CAPA. (B) *A. fumigatus* levels in those with and without CAPA measured by qPCR. (C) Beta diversity and CAPA status. (D) Relative abundance of *A. fumigatus* in mycobiomes of those with and without corticosteroid use. (E) *A. fumigatus* levels in those with and without corticosteroid use as measured by qPCR. (F) Dichotomised data for samples positive/negative for high *A. fumigatus* burden by qPCR in patients with or without corticosteroids. (G) Beta diversity and corticosteroid use. Boxplot data represent median and interquartile range. CS; corticosteroid use. CAPA; COVID-associated pulmonary aspergillosis.

**Fig. E4.**
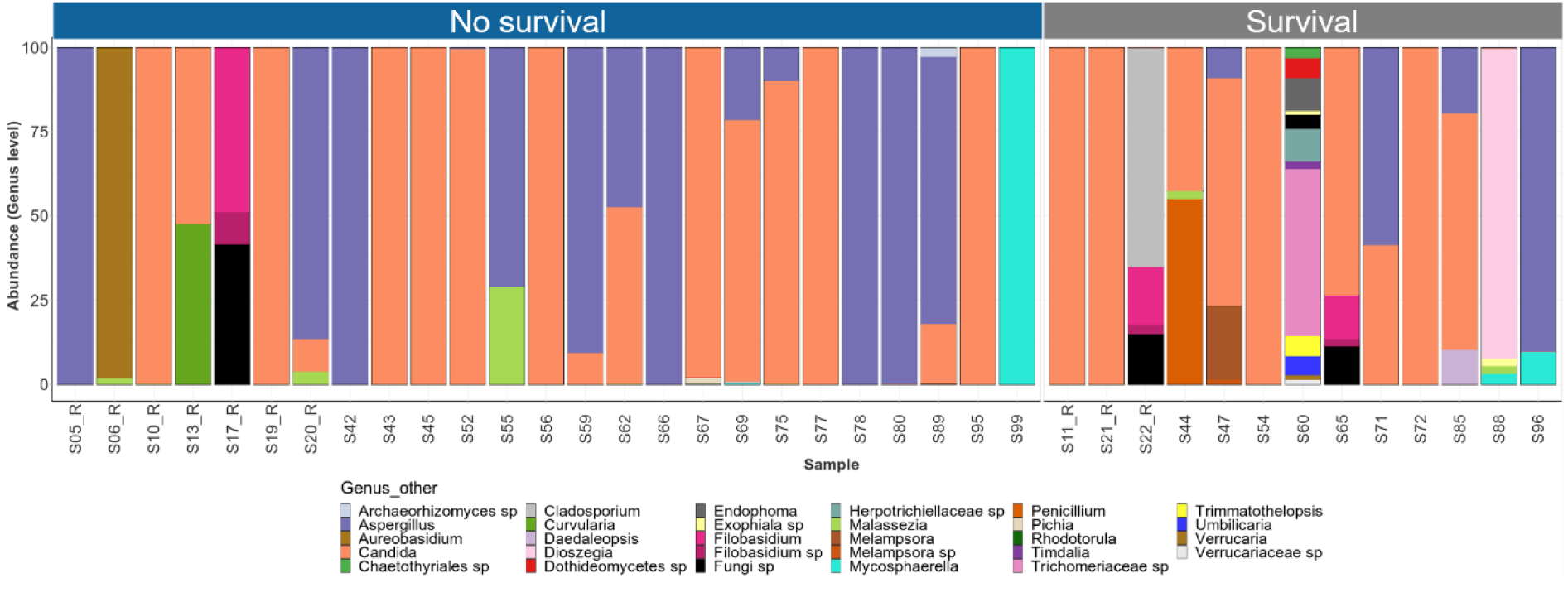
Sample data for ITS1 mycobiomes grouped by survival outcome. Relative abundance of fungal Genera identified in each sample grouped by survival outcome. Texa present below 1% were removed prior to visualisation.

**Fig. E5.**
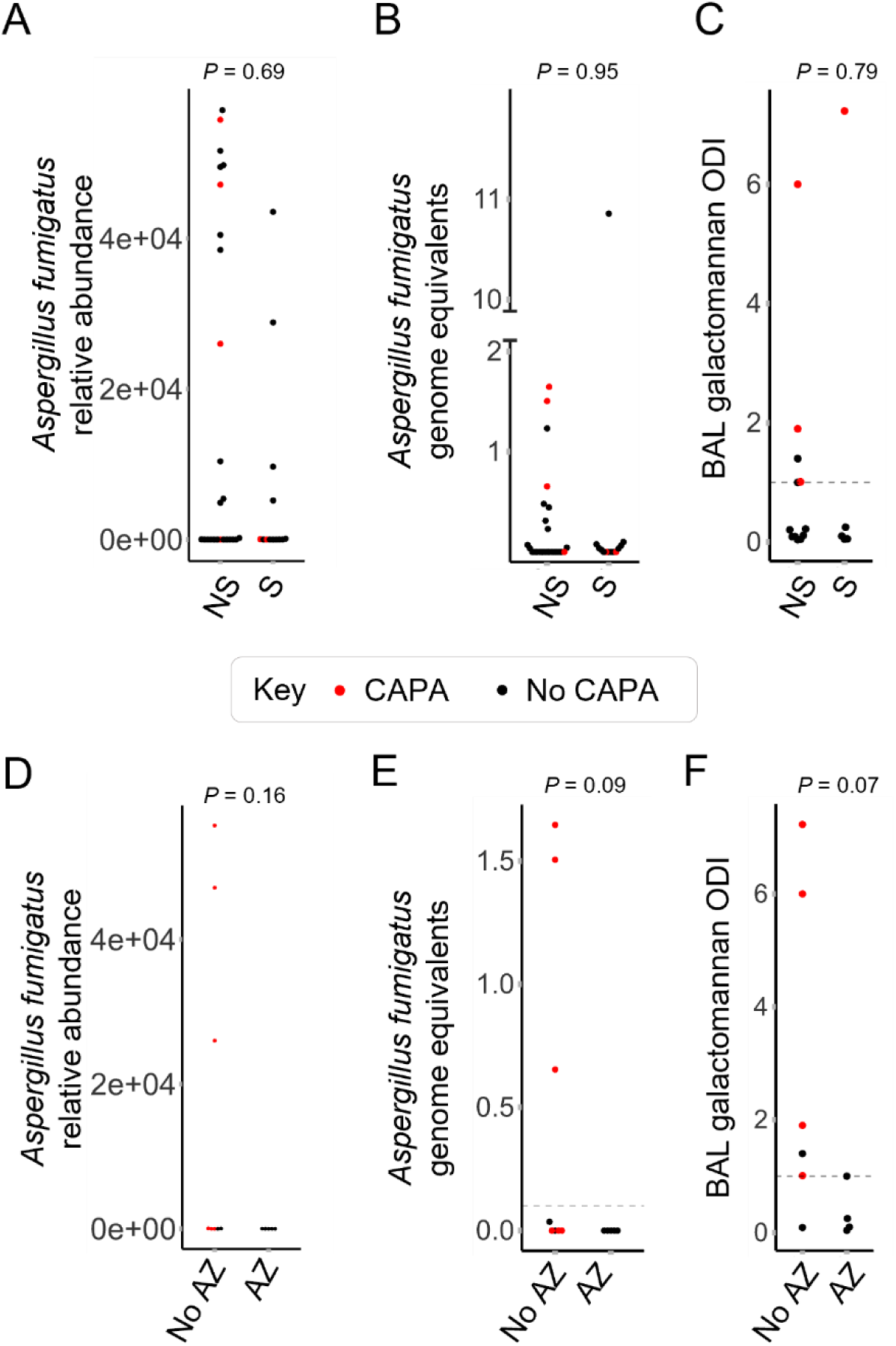
Extended data for *Aspergillus fumigatus* levels in respiratory mycobiomes and BAL galactomannan. (A) *Aspergillus fumigatus* relative abundance in patients grouped by survival outcome. (B) *A. fumigatus* burden measured by qPCR in patients which did not survive appeared biomodal, with some patients exhibiting a high burden (>0.1 genome equivalents). Excluding one outlier with an extremely high burden, all patients which survived displayed a low burden of 0.1 or below. (C) BAL galactomannan levels of patients grouped by survival outcome. (D) *Aspergillus fumigatus* relative abundance in patients grouped by azole treatment at ICU admission. (E) *A. fumigatus* burden measured by qPCR in patients which did or did not receive azole treatment at ICU admission. (F) All patients receiving azole treatment were BAL galactomannan negative (1 or lower). Half of patients not receiving azole treatment were galactomannan positive. CAPA: COVID-associated pulmonary aspergillosis. AZ: Azole treatment; No AZ: No azole treatment; S: Survival; NS: No survival.

**Fig. E6.**
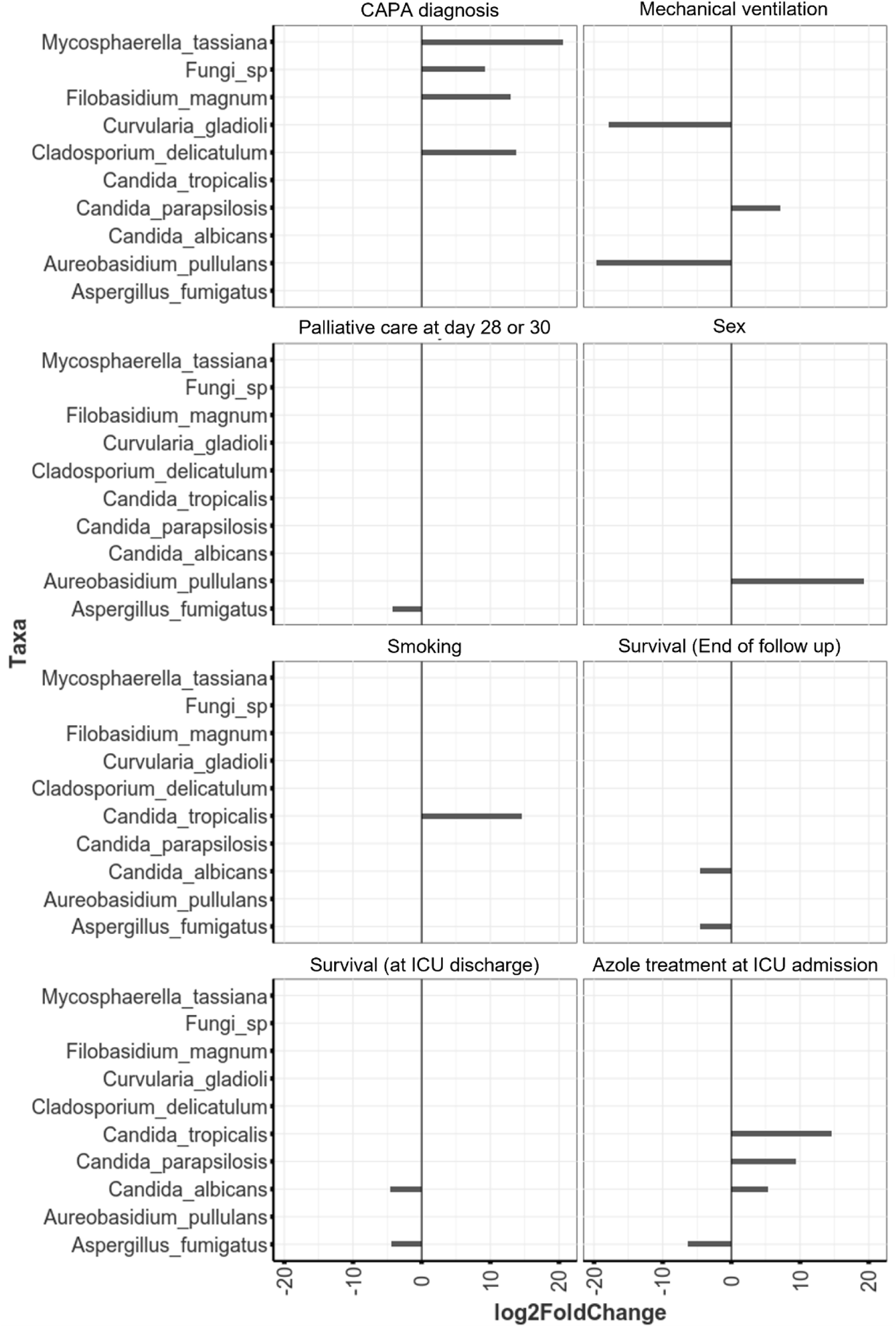
Significantly differentially abundant taxa identified by DESeq2 analysis.

**Table E1.** Full clinical patient demographics.

| Characteristics | N=39 |
| --- | --- |
| Age |  |
| Mean (SD) | 64 (7.4) |
| valid (missing) | 38 (1) |
| Sex |  |
| male | 82% (32) |
| female | 15% (6) |
| missing | 2.6% (1) |
| Ethnicity |  |
| Caucasian | 92% (36) |
| other | 5.1% (2) |
| missing | 2.6% (1) |
| BMI > 30 |  |
| yes | 23% (9) |
| no | 74% (29) |
| missing | 2.6% (1) |
| Smoking |  |
| yes | 15% (6) |
| no | 82% (32) |
| missing | 2.6% (1) |
| Institution |  |
| Graz | 26% (10) |
| Genoa | 62% (24) |
| Rennes | 13% (5) |
| Hematology oncology |  |
| yes | 10% (4) |
| no | 87% (34) |
| missing | 2.6% (1) |
| Solid organ transplant |  |
| yes | 5.1% (2) |
| no | 92% (36) |
| missing | 2.6% (1) |
| Cardiovascular disease |  |
| yes | 49% (19) |
| no | 49% (19) |
| missing | 2.6% (1) |
| Pulmonary disease |  |
| yes | 26% (10) |
| no | 72% (28) |
| missing | 2.6% (1) |
| Diabetes Mellitus |  |
| yes | 10% (4) |
| no | 87% (34) |
| missing | 2.6% (1) |
| Corticosteroids |  |
| yes | 74% (29) |
| no | 23% (9) |
| missing | 2.6% (1) |
| Tocilizumab |  |
| yes | 7.7% (3) |
| no | 90% (35) |
| missing | 2.6% (1) |
| Azithromycin |  |
| yes | 33% (13) |
| no | 62% (24) |
| missing | 5.1% (2) |
| Azole treatment at ICU admission |  |
| yes | 13% (5) |
| no | 21% (8) |
| missing | 67% (26) |
| Life Support |  |
| mechanical | 74% (29) |
| ECMO | 2.6% (1) |
| noninvasive | 5.1% (2) |
| mechanical & noninvasive | 10% (4) |
| none | 5.1% (2) |
| missing | 2.6% (1) |
| Duration ICU (days) |  |
| Mean (SD) | 31 (26) |
| valid (missing) | 38 (1) |
| Palliative (Day 28 or 32) |  |
| yes | 51% (20) |
| no | 46% (18) |
| missing | 2.6% (1) |
| Survival (at end of follow up) |  |
| yes | 33% (13) |
| no | 64% (25) |
| missing | 2.6% (1) |
| Antifungal Treatment Outcome |  |
| yes | 5.1% (2) |
| no | 5.1% (2) |
| partial | 0% (0) |
| missing | 90% (35) |
| Primary Treatment |  |
| yes | 5.1% (2) |
| no | 0% (0) |
| missing | 95% (37) |
| Antifungals initiated for CAPA |  |
| yes | 18% (7) |
| no | 0% (0) |
| missing | 82% (32) |
| Antifungal initiated |  |
| Posa Isa | 5.2% (2) |
| Isav | 0% (0) |
| IsavCasp | 2.6% (1) |
| LAmb | 0% (0) |
| Voriconazole | 10% (4) |
| missing | 82% (32) |
| BAL GM (ODI > 1) |  |
| positive | 21% (8) |
| negative | 72% (28) |
| missing | 7.7% (3) |
| BAL LFD |  |
| positive | 7.7% (3) |
| negative | 7.7% (3) |
| missing | 85% (33) |
| BAL PCR |  |
| positive | 7.7% (3) |
| negative | 49% (19) |
| missing | 44% (17) |
| BAL culture |  |
| positive | 7.7% (3) |
| negative | 90% (35) |
| missing | 2.6% (1) |
| Bronchial Aspirate culture |  |
| positive | 2.6% (1) |
| negative | 23% (9) |
| missing | 74% (29) |
| Tracheal Aspirate culture |  |
| Positive | 2.6% (1) |
| negative | 7.7% (3) |
| missing | 90% (35) |
| Tracheal Aspirate GM |  |
| positive | 2.6% (1) |
| negative | 2.6% (1) |
| missing | 95% (37) |
| Tracheal Aspirate PCR |  |
| positive | 5.1% (2) |
| negative | 21% (8) |
| missing | 74% (29) |
| Serum GM (> 0.5) |  |
| positive | 5.1% (2) |
| negative | 44% (17) |
| missing | 51% (20) |

**Table E2.** Full results for significantly differentially abundant taxa determined by DESeq2 analysis when assessing CAPA diagnosis, survival, and azole treatment at ICU admission.

| Variable | Taxa | base-Mean | log2FoldC hange | lfcSE | stat | p | Adjusted p |
| --- | --- | --- | --- | --- | --- | --- | --- |
| **CAPA diagnosis** | Cladosporium delicatulum | 4389.2 | 13.71 | 4.00 | 3.42 | 0.000619 | 0.003719 |
|  | Filobasidium magnum | 1165.8 | 12.93 | 3.13 | 4.13 | 3.59E-05 | 0.000368 |
|  | Fungi sp | 1017.7 | 9.18 | 2.62 | 3.51 | 0.000456 | 0.003719 |
|  | Mycosphaerella tassiana | 312608.4 | 20.62 | 4.01 | 5.15 | 2.65E-07 | 3.62E-06 |
| **Survival (at end of follow up)** | Aspergillus fumigatus | 8266.3 | -4.55 | 1.30 | -3.50 | 0.000462 | 0.017097 |
|  | Candida albicans | 83190.1 | -4.48 | 1.42 | -3.15 | 0.001611 | 0.029809 |
| **Azole treatment at ICU admission** | Aspergillus fumigatus | 943.1 | -6.33 | 2.43 | -2.6036 | 0.009225 | 0.042435 |
|  | Candida albicans | 66157.1 | 5.32 | 2.13 | 2.499698 | 0.01243 | 0.047648 |
|  | Candida parapsilosis | 8112.9 | 9.35 | 2.68 | 3.486701 | 0.000489 | 0.003749 |
|  | Candida tropicalis | 51931.1 | 14.52 | 2.99 | 4.85916 | 1.18E-06 | 1.36E-05 |

## Supplementary methods

### DNA extraction

BAL DNA was extracted using a cetyltrimethylammonium bromide (CTAB) method^24^. Negative extraction controls (NECs) consisting of molecular grade Tris-EDTA (TE) buffer (Promega) were included with each sample batch. First, 500 µl samples were incubated with 0.1% dithiothreitol (Thermo Scientific) at 37 °C for 45 minutes. To lyse fungal cells, CTAB DNA Extraction Buffer (Generon Ltd) was added and a short mechanical disruption (2 x 20 seconds FastPrep-24) using glass beads (425-600µM, Sigma) was performed, followed by 3 cycles of alternated heating to 65 °C and gentle vortexing for 10 mins each. Cell lysates were treated with 4 µl RNase A (100 mg/ml, Sigma) for 30 mins prior to DNA isolation using 1:1 ratio of Phenol-Chloroform (25:24:1, Merck) and 1:1 Chloroform:isoamyl alcohol (24:1, VWR). DNA was precipitated at -20 °C overnight using 2 volumes of ice-cold 100% ethanol and 0.1 volumes of 3M sodium acetate pH 4.8. Pelleted DNA was washed in 100% ethanol and twice in 70% ethanol. Final DNA was suspended in 50 µl TE buffer.

### qPCR

The presence of *A. fumigatus* in respiratory samples was validated using a TaqMan probe assay targeting the ITS1 region^26^. Assays were performed on an Applied Biosystems 7500 fast system using TaqMan Fast Advanced Master Mix (Thermo Fisher Scientific). Standard curves were a nine sample 10-fold dilution series beginning at 100 ng *A. fumigatus* genomes per reaction.

### PCR & Sequencing

The ITS1 region was amplified using Nextera XT compatible versions of ITS1^25^ and ITS2degen (a degenerate version of ITS2 primer) (see Table 1). PCR setup used Phusion Green Hot Start II HF PCR master mix to create 25 µl reactions with 300 nM forward primer, 1.2 µM reverse primer and 2 µl DNA. Parameters included annealing temperature of 55 °C for 40 cycles. PCRs included positive (*Aspergillus niger* genomic DNA), negative (molecular grade water) and negative extraction controls (NECs) from DNA extractions. Controls were processed fully and sequenced alongside the BAL samples. For sequencing library preparation, samples were processed as per Illumina fungal metagenomic demonstrated protocol (Starting at ‘Clean up’ section) using Nextera XT index kit v2 (Illumina). However, this protocol was modified to include sample pooling prior to the second clean up. Libraries were sequenced 2x150 on an Illumina iSeq100, with a final library loading concentration of 50 pM.

### Data analysis

Paired end reads were subject to quality trimming at Q30 and a minimum length filter of 75 nucleotides using bbduk^27^ (BBMap v38.22). Primer sequences were removed using Cutadapt^28^ (v1.18). Reads were mapped to UNITE database using bowtie2^29^ (v2.3.5.1). Count data was further processed in R (v4.1.3) using the following packages: phyloseq^30^ v1.38.0, vegan^31^ v2.5-7, DESeq2^32^ v1.34.0, stringr^40^ v1.4.0, ggplot2^41^ v3.3.5 and tidyr^35^ v1.2.0.

To minimise contamination in the mycobiome data, samples were subject to a multi-staged quality control process using NEC samples, along with qPCR and sequencing data. Firstly, samples from a DNA extraction batch with *A. fumigatus* detected (Ct <40) in the corresponding NEC by qPCR were excluded. Secondly, samples were excluded if they produced less than 500 fungal reads (read count filter) and/or a lower number of fungal reads than the corresponding NEC sample (NEC count filter). Of 99 initial samples, 22 were removed due to contamination of an extraction batch, 35 samples did not pass the read count filter and 3 did not pass the NEC count filter.

Abundances were standardised to the median sequencing depth. Extremely low abundance taxa were removed by only retaining those occurring > 0.2% in any sample. DESeq2 was used to identify significantly differentially abundant taxa (adjusted *p* value < 0.05 and basemean > 500). Differences in diversity (Shannon and observed OTUs) were assessed using pairwise Wilcoxon rank sum tests. PERMANOVA test was used to assess differences in Bray-Curtis ordination. A mycobiome sample was assigned as containing predominantly *A. fumigatus* if its *A. fumigatus* read counts were higher than 50% of the median sequencing depth. To determine mycobiome samples positive for high *A. fumigatus* burden by validatory qPCR, the cutoff was 0.1 haploid genome equivalents (HGE). The resulting contingency tables for sequencing and qPCR data were visualised as mosaic plots and statistical significance tested using Fisher’s exact tests.

## References

1. World Health Organisation. WHO Coronavirus Dashboard. https://covid19.who.int/.

2. Wiersinga, W. J., Rhodes, A., Cheng, A. C., Peacock, S. J. & Prescott, H. C. Pathophysiology, Transmission, Diagnosis, and Treatment of Coronavirus Disease 2019 (COVID-19): A Review. JAMA 324, 782–793 (2020).

3. Osuchowski, M. F. et al. The COVID-19 puzzle: deciphering pathophysiology and phenotypes of a new disease entity. Lancet. Respir. Med. 9, 622–642 (2021).

4. Dongelmans, D. A., et al. Characteristics and outcome of COVID-19 patients admitted to the ICU: a nationwide cohort study on the comparison between the first and the consecutive upsurges of the second wave of the COVID-19 pandemic in the Netherlands. Ann. Intensive Care 12, (2022).

5. Grasselli, G. et al. Hospital-Acquired Infections in Critically Ill Patients With COVID-19. Chest 160, 454–465 (2021).

6. Merenstein, C. et al. Signatures of COVID-19 Severity and Immune Response in the Respiratory Tract Microbiome. MBio 12, (2021).

7. Lloréns-Rico, V. et al. Clinical practices underlie COVID-19 patient respiratory microbiome composition and its interactions with the host. Nat. Commun. 2021 121 12, 1–12 (2021).

8. Ren, L. et al. Dynamics of the Upper Respiratory Tract Microbiota and Its Association with Mortality in COVID-19. Am. J. Respir. Crit. Care Med. 204, 1379–1390 (2021).

9. Gupta, A. et al. Mycobiome profiling of nasopharyngeal region of SARS-CoV-2 infected individuals. Microbes Infect. 105059 (2022) doi:10.1016/J.MICINF.2022.105059.

10. Ruiz-Rodriguez, A. et al. Bacterial and fungal communities in tracheal aspirates of intubated COVID-19 patients: a pilot study. Sci. Reports 2022 121 12, 1–10 (2022).

11. Kullberg, R. F. J. et al. Lung Microbiota of Critically Ill COVID-19 Patients are Associated with Non-Resolving Acute Respiratory Distress Syndrome. Am. J. Respir. Crit. Care Med. 1–74 (2022) doi:10.1164/rccm.202202-0274OC.

12. Viciani, E. et al. Critically ill patients with COVID-19 show lung fungal dysbiosis with reduced microbial diversity in patients colonized with Candida spp. Int. J. Infect. Dis. 117, 233–240 (2022).

13. Fraczek, M. G. et al. Corticosteroid treatment is associated with increased filamentous fungal burden in allergic fungal disease. J. Allergy Clin. Immunol. 1–8 (2017) doi:10.1016/j.jaci.2017.09.039.

14. Janssen, N. A. F. et al. Multinational Observational Cohort Study of COVID-19–Associated Pulmonary Aspergillosis. Emerg. Infect. Dis. 27, 2892 (2021).

15. Prattes, J. et al. Risk factors and outcome of pulmonary aspergillosis in critically ill coronavirus disease 2019 patients—a multinational observational study by the European Confederation of Medical Mycology. Clin. Microbiol. Infect. 28, 580–587 (2022).

16. Gangneux, J. P. et al. Fungal infections in mechanically ventilated patients with COVID-19 during the first wave: the French multicentre MYCOVID study. Lancet Respir. Med. 10, 180– 190 (2022).

17. Bartoletti, M. et al. Epidemiology of Invasive Pulmonary Aspergillosis Among Intubated Patients With COVID-19: A Prospective Study. Clin. Infect. Dis. 73, e3606–e3614 (2021).

18. Salmanton-García, J. et al. COVID-19–Associated Pulmonary Aspergillosis, March–August 2020 – Volume 27, Number 4—April 2021 – Emerging Infectious Diseases journal – CDC. Emerg. Infect. Dis. 27, 1077–1086 (2021).

19. Leistner, R. et al. Corticosteroids as risk factor for COVID-19-associated pulmonary aspergillosis in intensive care patients. Crit. Care 26, 1–11 (2022).

20. Koehler, P. et al. Defining and managing COVID-19-associated pulmonary aspergillosis: the 2020 ECMM/ISHAM consensus criteria for research and clinical guidance. Lancet Infect. Dis. 21, e149–e162 (2021).

21. Feys, S. et al. Lung epithelial and myeloid innate immunity in influenza-associated or COVID-19-associated pulmonary aspergillosis: an observational study. Lancet Respir. Med. 10, 1147– 1159 (2022).

22. Hoenigl, M. et al. COVID-19-associated fungal infections. Nat. Microbiol. 2022 78 7, 1127–1140 (2022).

23. Blot, S. I. et al. A clinical algorithm to diagnose invasive pulmonary aspergillosis in critically ill patients. Am. J. Respir. Crit. Care Med. 186, 56–64 (2012).

24. Fraczek, M. G. et al. The cdr1B efflux transporter is associated with non-cyp51a-mediated itraconazole resistance in Aspergillus fumigatus. J. Antimicrob. Chemother. 68, 1486–1496 (2013).

25. White, T. J., Bruns, T., Lee, S. & Taylor, J. Amplification and direct sequencing of fungal ribosomal RNA genes for phylogenetics. PCR Protoc. 315–322 (1990) doi:10.1016/B978-0-12-372180-8.50042-1.

26. Walsh, T. J. et al. Molecular Detection and Species-Specific Identification of Medically Important Aspergillus Species by Real-Time PCR in Experimental Invasive Pulmonary Aspergillosis. J. Clin. Microbiol. 49, 4150 (2011).

27. Bushnell, B. BBMap. sourceforge.net/projects/bbmap/ (2018).

28. Martin, M. Cutadapt removes adapter sequences from high-throughput sequencing reads. EMBnet.journal 17, 10 (2011).

29. Langmead, B. & Salzberg, S. L. Fast gapped-read alignment with Bowtie 2. Nat. Methods 9, 357–359 (2012).

30. McMurdie, P. J. & Holmes, S. phyloseq: An R Package for Reproducible Interactive Analysis and Graphics of Microbiome Census Data. PLoS One 8, e61217 (2013).

31. Dixon, P. VEGAN, a package of R functions for community ecology. J. Veg. Sci. 14, 927–930 (2003).

32. Love, M. I., Huber, W. & Anders, S. Moderated estimation of fold change and dispersion for RNA-seq data with DESeq2. Genome Biol. 15, 1–21 (2014).

33. Wickham, H. stringr: Simple, Consistent Wrappers for Common String Operations. R package version 1.4.0. https://CRAN.R-project.or at (2019).

34. Wickham, H. ggplot2: Elegant Graphics for Data Analysis. at (2019).

35. Wickham, H. & Henry, L. tidyr: Tidy Messy Data. R package version 1.1.0. https://CRAN.R-project.or at (2020).

36. Nguyen, L. D. N., Viscogliosi, E. & Delhaes, L. The lung mycobiome: an emerging field of the human respiratory microbiome. Front. Microbiol. 6, 89 (2015).

37. Huang, C. et al. Fungal and bacterial microbiome dysbiosis and imbalance of trans-kingdom network in asthma. Clin. Transl. Allergy 10, 1–13 (2020).

38. Martinsen, E. M. H. et al. The pulmonary mycobiome-A study of subjects with and without chronic obstructive pulmonary disease. PLoS One 16, 1–16 (2021).

39. Tiew, P. Y. et al. A high-risk airway mycobiome is associated with frequent exacerbation and mortality in COPD. Eur. Respir. J. 57, (2021).

40. Wickham, H. stringr: Simple, Consistent Wrappers for Common String Operations. at https://cran.r-project.org/package=stringr (2019).

41. Wickham, H. ggplot2: Elegant Graphics for Data Analysis. *Springer-Verlag New York* (2016).

